# Mechanistic analysis of compounds that modulate VEGF-A splicing in podocytes with therapeutic potential for diabetic nephropathy

**DOI:** 10.1101/2025.11.27.690981

**Authors:** Monica Ayine, Megan Stevens, Sebastian Oltean

## Abstract

Vascular endothelial growth factor A (VEGF-A) has an alternatively spliced variant, VEGF-A165b, formed when a distal 3’ splice site in exon 8 is selected. The anti-angiogenic and anti-permeability VEGF-A165b has reno-protective properties and has been shown to rescue kidney function in diabetic nephropathy (DN) mouse models (PMID 25542969). We previously identified three new compounds that can increase the expression of the VEGF-A165b isoform. Two of these compounds, named ESSO1 and ESSO3, were discovered through a screen of synthetic compounds that alter VEGF-A splicing (PMID 33941763). A third compound, delphinidin, was found to be the key regulator of VEGF-A splicing in a natural blueberry and sea-buckthorn extract (DIAVIT) (PMID 30865689).

This study aimed to investigate the signalling mechanism through which these three compounds regulate VEGF-A splicing and increase the protective VEGF-A165b isoform in renal podocytes.

Human podocytes were exposed to a diabetic environment (glucose soup [GS]: 25 mM glucose, 1 ng/ml TNF-_α_, 1 ng/ml IL-6, and 100 nM insulin), in comparison to a normal glucose (5.5 mM glucose) and an osmotic control (5.5 mM glucose + 19.5 mM mannitol) control, for 48 hours. RNA and protein were extracted for RT-PCR and Western blotting analysis of splice isoforms. Immunofluorescence and cell viability assays were also used.

Immunofluorescence showed no significant difference in nephrin expression for all treatments compared to control, suggesting that normal podocyte function is not affected by these compounds. Trypan blue exclusion assay, used to assess the percentage cell viability of podocyte cells, showed that they do not affect podocyte cell viability.

By using various inhibitor treatments, we have found several molecules involved in signalling, like JNK and splicing kinases SRPK1 and CLK1 for ESSO1, splicing factor hnRNPH and kinase CLK1 for ESSO3 and AMPK, splicing factor SRSF6 and kinase CLK1 for delphinidin. Other molecules, like ATM, E2F1, and hnRNPA1, were ruled out.

Interestingly, all three compounds significantly increased expression of the splicing kinase CLK1, which has been involved in VEGF-A splicing regulation, suggesting that all three compounds may be increasing the VEGF-A_165_b/panVEGF-A_165_ ratio through signalling through this kinase. Studies of human biopsies showed that CLK1 is downregulated in DN patients compared to healthy volunteers. The development of a CLK1 activator as a DN therapy seems, therefore, a plausible opportunity.

## 1. Introduction

Alternative splicing (AS) is a highly regulated post-transcriptional process that gives rise to multiple proteins from the same gene via exon skipping/inclusion or intron inclusion (1). Given that more than 94% of human genes are estimated to undergo AS, it is a key source of proteome diversity (2,3). In some cases, AS isoforms display functionally opposite properties and behave as though they were transcribed from different genes (4).

AS is predominantly carried out by the spliceosome, a large ribonucleoprotein complex. The spliceosome catalyses two transesterification reactions, removing introns and joining exons. To recognise sequences that must be removed from a transcript to form the mature mRNA, cis-acting elements within the pre-mRNA and trans-acting factors, which are primarily RNA-binding proteins (RBPs), act together to direct the spliceosome(1).

The vascular endothelial growth factor A (*VEGF-A*) gene has eight exons and seven introns. The use of the proximal splice site (PSS) in exon 8 gives rise to several pro-angiogenic variants collectively known as VEGF-A_xxx_a, where xxx indicates the number of amino acids in the polypeptide chain. A novel 3’ distal splice site (DSS) was later discovered in exon 8, 66 base pairs downstream of the PSS splice site (5,6). When the DSS is selected, a functionally different anti-angiogenic isoform is formed, termed VEGF-A_xxx_b. VEGF-A_xxx_a and its VEGF-A_xxx_b counterparts have the same number of amino acids; however, they differ in the six C-terminal amino acid sequence, giving the VEGF-A_xxx_b isoform an opposite functionality to that of VEGF-A_xxx_a (7). The most predominant VEGF-Axxxb isoform, VEGF-A165b, has been shown to bind to VEGF receptor-2 (VEGFR-2) with the same affinity as its canonical form; however, the phosphorylation and subsequent activation of VEGFR-2 are reduced (Woolard et al., 2004) (6,8). Due to the contrasting functions of these isoforms, an increase in the expression of one isoform over the other may cause an increase or repression in angiogenesis. It has been hypothesised that the reason angiogenesis is not detected in the renal cortex is due to the balance in the expression of the pro- and anti-angiogenic isoforms (9). Although there is evidence that points to an imbalance in the isoforms in diseases such as diabetic nephropathy (DN), where the expression of the anti-angiogenic VEGF-A_165_b decreases as the disease progresses, the molecular pathways that regulate the AS of exon 8 in the *VEGF-A* gene have not yet been fully elucidated. Given that the recombinant VEGF_165_b protein has been shown to have therapeutic potential in models of DN (10) it is important to understand how compounds that modulate *VEGF-A* splicing work, with the potential to develop them in therapeutics in the future.

The expression of a particular protein isoform is dependent upon splice site selection. The recognition of splice sites by the spliceosome relies on RBPs, mainly SR proteins and heterogeneous nuclear ribonucleoproteins (hnRNPs), binding to auxiliary sequences, such as exonic splicing enhancers (ESEs) and intronic splicing enhancers (ISEs) found in nascent transcripts. The SR-protein SRSF1 has been shown to regulate AS of the *VEGF-A* gene in favour of the PSS to produce VEGF-A_xxx_a (7,11). In contrast, SRSF2 and SRSF6 influence the selection of the DSS (**Supplementary Figure**) (12,13). To mediate splice site selection, SR-proteins require activation through phosphorylation. SRPK1 and CLK1 are the most common kinases that phosphorylate SR-proteins involved in *VEGF-A* AS (**Supplementary Figure**). However, these kinases are also regulated by other factors, such as extracellular signalling. In addition, growth factors, including IGF-1 and TNF_α_, can stimulate VEGF-A_xxx_a expression, whereas TGF-β1 promotes the expression of VEGF-A_xxx_b (7).

In a previous paper, we performed a screen that identified a number of compounds that could modulate *VEGF-A* splicing to promote VEGF-A_xxx_b, with potential to be developed as therapeutics (14). Two of the most promising compounds were called ESSO1 and ESSO3. (15)ESSO1 is trovafloxacin; it is a broad-spectrum antibiotic and a fourth-generation fluoroquinolone. It is more effective against gram-positive than gram-negative bacteria, and the most potent fluoroquinolone against anaerobic bacteria. Its antibacterial activity functions through DNA gyrase and topoisomerase IV inhibition (16). Trovafloxacin was withdrawn from the market due to hepatotoxicity concerns (17); however, its use is now reserved for life or limb-threatening infections. ESSO3 (5-[(4-Ethylphenyl) methylene]-2-thioxo-4-thiazolidinone) is a c-MYC inhibitor that induces cell cycle arrest and apoptosis (18). It specifically inhibits the dimerisation of c-MYC–MAX. C-MYC is a well-studied transcription factor due to its role as a proto-oncogene.

An additional paper (15) showed that Delphinidin, a natural pigment belonging to the anthocyanidin family, can also modulate *VEGF-A* splicing to promote VEGF-A_xxx_b. Delphinidin is commonly found in berries and gives a blue-red colour to flowers and fruits. It is known to have antioxidant and anti-inflammatory properties and is involved in many biological processes, such as apoptosis, autophagy, and angiogenesis (19).

To understand how ESSO1, ESSO3, and delphinidin modulate AS of the *VEGF-A* gene, we need to understand the signalling pathways associated with each compound. Furthermore, it is important to understand whether the pathways by which they modulate *VEGF-A* splicing in podocytes is similar to those reported in other cell systems. Therefore, this study aimed to elucidate the pathways through which these three compounds modulate the AS of *VEGF-A*.

## 2. Materials and Methods

### 2.1 Cell culture and treatment

The human renal epithelial cell line Podo/TERT256 (Evercyte) and human embryonic kidney cell line HEK293 were sub-cultured from existing cultures in the lab at 37 °C, 5% CO_2_, and 90–95% humidity. Podo/TERT256 cells were cultured in EGM-2 media (Lonza; CC-3162) supplemented with the EGM-2 bullet kit containing foetal bovine serum (FBS), growth factors, and antibiotics (Lonza; CC-3162), without VEGF. HEK293 cells were cultured in low glucose DMEM ( Sigma–Aldrich; D6046) supplemented with 10% FBS (GIBCO; 10270106) and 1% penicillin–streptomycin (Sigma-Aldrich; P4333).

Cells were treated in a diabetic environment and were compared to cells treated in normal glucose (NG), as well as a mannitol (M) osmotic control treatment, for 48 h. The treatment media contained no growth factors and 0.5% FBS (Sigma–Aldrich; F7524).

Prolonged in vitro culture of kidney cells can lead to degradation of insulin receptors. Therefore, to mimic a diabetic environment and induce cellular insulin resistance, the cells were exposed to a concentrated “glucose soup” treatment(20,21). The NG treatment contained 5 mM glucose; the diabetic environment treatment, termed glucose soup (GS), consisted of 25 mM glucose, 1 ng/ml TNFα, 1 ng/ml IL-6, and 100 nM insulin; and the M treatment contained 5 mM glucose and 20 mM mannitol. Other treatments, such as the small molecules under investigation, were added to the treatment media. Cells were treated with ESSO(1-9) at a concentration of 10 μM and Delphinidin at a concentration of 10 μg/ml and compared with vehicle controls. SPHINX was used as a positive control to assess *VEGF-A* splicing(22).

### 2.2 Growth curve

Cells were seeded at a density of 100,000 cells per well in four 12-well plates and treated 24 h later. The cells were detached using trypsin, resuspended in a serum-free medium, and counted using a counting chamber (Marienfeld). Samples were counted every 24 h, and the treatment media was refreshed 48 h after the first treatment.

### 2.3 Alamar blue assay

Alamar blue is a resazurin-based cell viability reagent. Resazurin is an oxidation-reduction (REDOX) indicator that changes colour through an oxidation-reduction reaction. Alamar blue is non-toxic and permeable to cells. On entering living cells, the blue non-fluorescent resazurin changes to its reduced form, called resorufin, which is pink and highly fluorescent. The fluorescent intensity measured corresponds to the number of respiring living cells. Following cell treatment for 48 h, the media was replaced with serum-free media containing 1% Alamar Blue and incubated for 3 h. A bottom fluorescence reading of each well was taken at an excitation of 560 nm and an emission of 590 nm using a Plate Reader (Molecular Devices SupraMax M2). Percentage viability was calculated as follows (FI, fluorescence intensity): (FI 590 treated cells / FI 590 control cells) × 100.

### 2.4 Wound healing/ scratch assay

Samples were cultured to form a monolayer of cells at 70–80% confluency. Two horizontal lines were made across the bottom of the well, creating a scratch in the monolayer. The media was replaced, and photomicrographs of the initial gap area at 0 h and the residual gap area every 24 h until 48 h were taken. The wound area was calculated from photomicrographs and analysed by ImageJ.

### 2.5 Trypan Blue assay

The trypan viability assay is based on the exclusion of the dye from live cells. Trypan blue is impermeable to the cell membrane; therefore, it only enters compromised cells and binds to intracellular proteins. As it concentrates in cells that are not viable, it gives off a dark-blueish colour.

The cells were detached using trypsin, and fresh medium with treatment compounds was added. An equal volume of re-suspended cells and filtered trypan blue was mixed and incubated for 2–3 min at room temperature. The percentage of cell viability was assessed using the Bio-Rad TC20 Automated Cell Counter.

### 2.6 RT-qPCR

RNA was extracted from cells using the RNeasy kit (Qiagen) or the phenol–chloroform extraction method before reverse transcribing to cDNA(23). Real time quantitative polymerase chain reaction (RT-qPCR) was performed with target primers (**Table 1**) using SYBR green (Applied Biosystems A25742). The cycling conditions were as follows: 95 °C for 60 s, followed by 45cycles of 95 °C 15 s, 58 °C for 15 s and 72 °C for 60 s.

### 2.7 Western Blot

Protein was extracted from cells using RIPA lysis buffer (Thermos Scientific) with protease and phosphatases inhibitor cocktail (Thermos Scientific). Protein samples underwent sodium dodecyl sulfate–polyacrylamide gel electrophoresis (SDS–PAGE) with 4–15 % Mini-PROTEAN TGX stain-free precast polyacrylamide gels (BIORAD). The gel was imaged to activate the TGX, which allows for visualisation and accurate analysis of the total protein loaded for each sample using a Gel-Doc EZ (BIO-RAD) imaging system. The use of this system means a housekeeping protein loading control is not required as the amount of protein on the membrane for each sample can be quantified. Following protein transfer onto a polyvinylidene fluoride (PVDF) membrane (ThermoFisher), blocking was performed with 3% bovine serum albumin (BSA, Sigma–Aldrich) in tris-buffered saline (TBS)-Tween (0.1%) for 1 h at room temperature. The membranes were subsequently probed with primary anti-mouse hVEGF_165_b and anti-rabbit panVEGF (Abcam) at a 1:1000 dilution in 3% BSA TBS-Tween, overnight at 4°C with agitation. The membranes were washed and incubated in secondary antibody at a 1:10000 dilution in 3% BSA TBS-Tween for 2–3 h at room temperature in the dark with agitation. After washing membranes in TBS-Tween (0.3%), they were incubated in fluorescent secondary antibodies (LI-COR) diluted in 3% BSA-TBS-Tween (0.3%), 1:10,000. Membranes were washed again and imaged with the LI-COR Odyssey® CLx. Analysis was performed using the Image Studio software (LI-COR).

### 2.8 Immunofluorescence (IF)

The cells were seeded at a density of 15,000 cells per ml onto cover slips placed in a 12 well plate. Following compound treatment, the cells were washed in ice-cold phosphate-buffered saline (PBS) and incubated in ice-cold 100% methanol for 5 min to fix the cells. Cover slips were washed in ice-cold PBS and incubated in 1% BSA in PBS with 0.1% Tween (Sigma-Aldrich) for 30 min to prevent nonspecific antibody binding. The cell were then probed with a primary antibody, nephrin (B-12) conjugated with ALEXA Fluor 488 (Santa Cruz Biotechnology sc-377246), diluted at 1:50–1:250 in blocking solution and incubated for 90 min at room temperature in the dark. Finally, the coverslips were washed and placed face down on a microscope slide with a small drop of Fluoroshield. The edges were carefully sealed and then imaged.

### 2.9 Immunoprecipitation

Cells were seeded at a density of 20,000 cells per ml into a 6 well plate 12 h before serum starvation for 4–6 h. Following serum starvation, cells were treated for 2 h in serum-free media and re-treated with or without EGF at 25 ng/ml for 1 h. Samples were washed with ice-cold PBS with protease and phosphatase inhibitors, lysed, collected in a tube and incubated on ice for 15 min. The tubes were centrifuged at 10,000 rpm for 15 min at 4°C and the supernatant was transferred to a fresh tube on ice. Anti-SR (1H4) was added at a 1:10 dilution and incubated on a tube spinner at 4°C overnight. Pierce protein A/G magnetic beads (ThermoScientific) were added to each tube and incubated on a rocker for 2 h at 4°C. Tubes were centrifuged at 10,000 rpm for 5 min, the supernatant discarded, and the pellet washed with RIPA and a protease and phosphatase inhibitor cocktail five times. The pellet was then re-suspended, vortexed thoroughly, and beta mercaptoethanol (BIORAD)loading dye was added followed by heating to 95°C for 5 min. The samples underwent Western blotting with a mouse mAb104 primary antibody (anti-phospho-SR) diluted at 1:4 dilution in TBS-Tween(24).

### 2.10 JC-10 assay

The JC-10 assay detects changes in the mitochondrial membrane potential using the cationic, lipophilic JC-10 dye. In normal cells, JC-10 concentrates in the mitochondrial matrix where it forms red fluorescent aggregates. However, in apoptotic and necrotic cells, JC-10 diffuses out of the mitochondria, changes to monomeric form, and stains cells with green fluorescence. We display the data as the mitochondrial polarisation relative to control as a marker of apoptosis. Cells were plated at a density of 20,000 per well in 90 μl media in a clear-bottom black 96-well cell culture plate and left to adhere overnight. Following treatment for the desired time, 50 μL of the JC-10 dye-loading solution, consisting of Assay Buffer A and the JC-10 probe (Abcam) was added to each well, per the manufacturer’s instructions. The plate was then incubated for 1 h, protected from light. After incubation, 50 μL of Assay Buffer B was added to each well, and the FI was measured using a fluorescent plate reader. Excitation/emission at 490/525 nm, with cut off at 515 nm, measured green fluorescence, while 540/590 nm (with cut off at 570 nm) measured red fluorescence. The green/red FI ratio was calculated.

### 2.11 Statistical analysis

Data were analysed using GraphPad Prism (Prism 9). The data were first tested for normality and lognormality. An unpaired student t-test (parametric) or Mann–Whitney t test (nonparametric tests) was used to compare a two-group data set, and for three or more groups of datasets, a one-way analysis of variance (ANOVA) parametric, lognormal ANOVA or Kruskal–Wallis (nonparametric tests) was used. Results were considered statistically significant if the P value was < 0.05.

## 3. Results

### 3.1 ESSO1, ESSO3, and delphinidin consistently increase VEGF-A_165_b levels in podocytes

In a previous screen (14), we identified nine compounds (**Table 1**) that could switch *VEGF-A* splicing to increase VEGF-A_xxx_b isoforms. The studies were performed in human kidney embryonic (HEK293) cells and prostate cancer (PC3) cells. Since the regulation of AS is cell-specific, we wanted to test which compounds consistently increase VEGF-A_165_b in podocytes. In Podo/TERT256 cells, treatment with ESSO1 and EESO3 resulted in a significant increase in the protein VEGF-A_165_b/panVEGF-A_165_ ratio after 48 h (**Figure 1 A, B**). Therefore, we decided to focus the remainder of this study on the therapeutic potential of ESSO1 and ESSO3 in DN.

**Figure 1.**
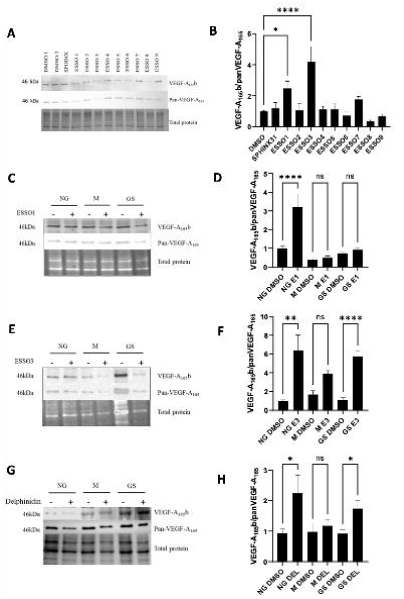
Western blots and quantifications showing VEGF_165_b expression levels upon treatment of podocytes with ESSOs or Delphinidine in normal or diabetic conditions. **A.** Podocyte cells were treated at a concentration of 10 μM of the nine ESSOs for 48 hrs in growth factor-free and serum-reduced media. Image of a Western blot performed after proteins were extracted from treated cells **B.** Quantification of the Western blot showing a significant increase in VEGF-A_165_b with respect to panVEGF-A_165_ when podocyte cells were treated with ESSO1 (mean=2.48, ±0.45 SEM, *p=0.0212, N=17 repeats) and ESSO3 (mean= 4.20, ± 0.97 SEM, ****p<0.0001, N=18 repeats), compared to control **C.** Podocyte cells were treated for 48 hrs at 10 μM with ESSO1 under normal glucose (NG: 5 mM glucose), high glucose soup conditions (GS: 25 mM glucose, 1 ng/ml TNFα, 1 ng/ml IL-6, 100 nM insulin), mannitol osmotic control (M: 5 mM glucose + 20 mM mannitol) and a vehicle control (DMSO), all in growth factor free and serum reduced media. Image of a Western blot performed after proteins were extracted from treated cells **D.** Quantification of the Western blot showing a significant increase in VEGF-A_165_b with respect to panVEGF-A_165_ in normal glucose environment (mean= 3.21, ± 0.69 SEM, ****p<0.0001, N=9 repeats) compared its DMSO control, however, VEGF-A_165_b levels were notable low under diabetic conditions. **E.** Podocyte cells were treated for 48 hrs at 10 μM with ESSO3 under normal glucose (NG: 5 mM glucose), high glucose soup conditions (GS: 25 mM glucose, 1 ng/ml TNFα, 1 ng/ml IL-6, 100 nM insulin), mannitol osmotic control (M: 5 mM glucose + 20 mM mannitol) and a vehicle control (DMSO), all in growth factor free and serum reduced media. Image of a Western blot performed after proteins were extracted from treated cells **F.** Quantification of the Western blot showing a significant increase in VEGF-A_165_b with respect to panVEGF-A_165_ in normal glucose environment (mean= 6.39, ± 1.66 SEM, **p<0.0058, N=9 repeats) compared its DMSO control, as well as a significant increase in VEGF-A_165_b levels in diabetic conditions (mean= 5.75, ± 0.53 SEM, ****p<0.0001, N=9 repeats). **G.** Podocyte cells were treated for 48 hrs at 10 ug/ml with Delphinidin under normal glucose (NG: 5 mM glucose), high glucose soup conditions (GS: 25 mM glucose, 1 ng/ml TNFα, 1 ng/ml IL-6, 100 nM insulin), mannitol osmotic control (M: 5 mM glucose + 20 mM mannitol) and a vehicle control (DMSO), all in growth factor free and serum reduced media. Image of a Western blot performed after proteins were extracted from treated cells **H.** Quantification of the Western blot showing a significant increase in VEGF-A_165_b with respect to panVEGF-A_165_ in normal glucose environment (mean= 0.5951, ± 1.49 SEM, *p=0.0234, N=8 repeats) compared its DMSO control as well as a significant increase in VEGF-A_165_b levels in diabetic conditions (mean= 1.740, ± 0.26 SEM, *p<0.0455, N=8 repeats).

The effect of ESSO1 and ESSO3 on VEGF-A splicing in Podo/TERT256 cells under diabetic conditions (GS) was examined using Western blotting. Consistent with the preliminary results, both ESSO1 and ESSO3 increased the VEGF-A_165_b/panVEGF-A_165_ (**Figure 1C–F**) ratio in NG conditions after 48 h. While an increase of VEGF-A_165_b/panVEGF-A_165_ ratio increased with ESSO3 treatment under diabetic conditions (**Figure 1E, F**), ESSO1 treatment did not cause the same increase under diabetic conditions (**Figure 1C, D**).

Delphinidin was previously reported to increase VEGF-A_165_b expression (15). In Podo/TERT256 cells, we found that delphinidin increased the VEGF-A_165_b/panVEGF-A_165_ ratio under NG conditions after 48 h. There was also significant increase in VEGF-A^165^b/panVEGF-A^165^ under diabetic conditions **(Figure 1G,H)**.

### 3.2 Effect of ESSO1, ESSO3, and delphinidin on various cellular properties

We further wanted to characterise the effect of the compounds on podocyte-specific markers, such as nephrin, as well as their effects on cellular viability, growth, migration, and apoptosis, to obtain a more in-depth understanding of how these compounds affect podocytes.

#### Podocyte nephrin expression is t affected by ESSO3,but not ESSO1 or delphinidin

To elucidate the impact of the three compounds on the podocyte marker nephrin, IF using a nephrin-specific antibody was performed on Podo/TERT256 cells treated with the compounds for 48 h. We observed a non-significant increase in nephrin expression when podocyte cells were treated with ESSO1 and delphinidin compared to the DMSO control treatment. However, ESSO3 treatment showed a significant decrease in nephrin expression compared to the DMSO control (**Figure 2A, B**).

**Figure 2.**
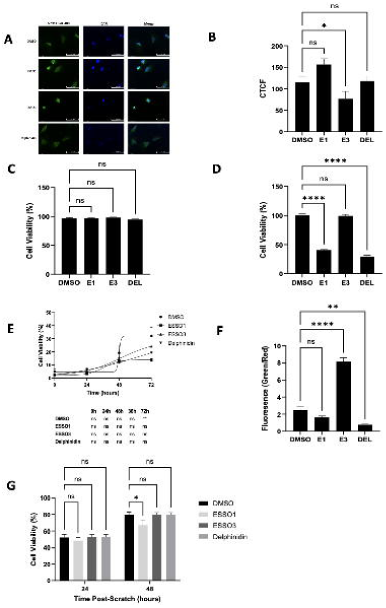
The effect of ESSO 1, ESSO3 and Delphinidin on various cellular properties. **A.** Immunofluorescence staining images of podocyte cells treated with either DMSO, ESSO1 (10 μM), ESSO3 (10 μM) or Delphinidin (10 μg/ml) for 48 hrs. Cells were stained for nephrin using a nephrin antibody coupled with Alexa Fluor® 488 (in green) and the nuclei were stained with DAPI (in blue). **B.** Fluorescence quantification for each treatment. Magnification: 40X, (p=ns; N=9 repeats). Data analyses was performed using lognormal ordinary one way ANOVA analysis. ESSO3 reduced nephrin expression significantly (mean= 76.53, ± 17,12 SEM, *p=0.031, N=9 repeats). **C.** Trypan Blue cell viability assay shows no significant difference between the treatments (ESSO1, ESSO3 or Delphinidin) and DMSO control (p=ns; N=12 repeats). Data were analysed using a non-parametric Kruskal-Wallis post-hoc analysis. **D.** Alamar blue assay was performed for HEK293 cells treated with for 48 hrs with the three compounds. Fluorescence was measured by at 560nm excitation /590nm emission 3 hrs after adding the reagent. ESSO1 and Delphinidin both significantly reduced cell viability, ****p<0.0001, whilst ESSO3 had no effect ns p=0.9974 (N=16 repeats). Error bars represent SEM. Data were analysed by lognormal Brown-Forsythe and Welch ANOVA test. **E.** HEK293 cells treated with the three compounds were counted at 24 hrs and then every 24 hrs until 72 hrs when growth plateaued. At 72 hrs, ESSO1 significantly inhibit cell growth (**p<0.0048). Both ESSO3 and Delphinidin did not significantly affect HEK293 cell growth. Data were analysed using a two-way ANOVA (N=6 repeats). **F.** Cells were plated and treated for 48 hrs. A plate reader was used to assess the red/green fluorescence. The graph represents a ratio of apoptotic cells (green) to normal cells (red). ESSO3 induced apoptosis (****p<0.0001), whilst Delphinidin was the only compound to inhibit apoptosis (**p<0.0013), ESSO1 did not affect apoptosis (p=0.1878). Error bars represent SEM. Data were analysed by ordinary one-way ANOVA (N=12 repeats). **G.** After creating a gap in confluent monolayer cells, they were treated, and images of the gap were taken at 24 and 48h. Analysis of the rate of gap closure was done with image J, ESSO1 inhibited cell migration significantly after 48 hrs (*p=0.0387). Error bars represent SEM. Data were analysed by ordinary one-way ANOVA (N=3 repeats).

#### Trypan Blue

The trypan blue assay was used to assess the effects of the compounds on podocyte cell viability. We observed no significant difference in viability following treatment with all three compound treatments for 48 h compared to the DMSO control (**Figure 2C**).

#### Alamar Blue

The Alamar blue assay was also used to assess cell viability. We observed a significant reduction in cell viability after HEK293 cells were treated with ESSO1 and delphinidin compared to the DMSO control. ESSO3, however, did not show any significant change in cell viability compared to the DMSO control (**Figure 2D**).

#### Growth curve

Growth curves were generated by plating an equal number of HEK293 cells at time zero and then counting cells after 24, 48, and 72 h. The compounds did not impact cell proliferation in comparison to the DMSO control after 24 h. However, after 48 h, cell proliferation plateaued with ESSO1 treatment, with a significant decrease compared to the DMSO control. ESSO3 and delphinidin also reduced proliferation; however, this reduction was not significant compared to the DMSO control (**Figure 2E**).

#### Apoptosis

The JC-10 assay was used to determine whether the compounds induced apoptosis in HEK293 cells. The ratio of green/red is proportional to the number of apoptotic cells. The assay showed a significant increase in HEK293 apoptosis following treatment with ESSO3 compared to the DMSO control. In contrast, delphinidin resulted in a significant decrease in apoptosis. ESSO1, however, did not cause any change in apoptosis (**Figure 2F**).

#### Wound healing/scratch assay

To examine the influence of the compounds on cell migration, the scratch or wound healing assay was employed. The rate of wound closure was measured at different time points. ESSO1 had a small inhibitory effect on cell migration after 48 h compared to the DMSO control. In contrast, ESSO3 and delphinidin did not affect cell migration (**Figure 2G**).

### 3.3 Effects of ESSO1, ESSO3, and delphinidin on the expression of PSS selection regulators, SRPK1 and SRSF1, and DSS selection regulators, CLK1 and SRSF6

To enable the further development of these compounds as diabetic kidney therapeutics, we investigated their mechanism of action, aiming to describe how they signal to the RNA and spliceosome to change splicing. We initially based the investigation on molecules that have been characterised previously (not necessarily in kidney cells) to regulate VEGF-A terminal exon splicing (7) (**Supplementary Figure**).

The best-characterised molecules that drive PSS selection in VEGF-A exon 8 are the splicing kinase SRPK1 and its substrate SRSF1. Therefore, we assessed whether the compounds change the expression of these mRNAs in podocytes after 48 h. As shown in **Figures 3A and B**, none of the three compounds had an effect on either SRPK1 or SRSF1 mRNA expression, with the exception of ESSO3, which significantly reduced SRSF1 levels.

**Figure 3.**
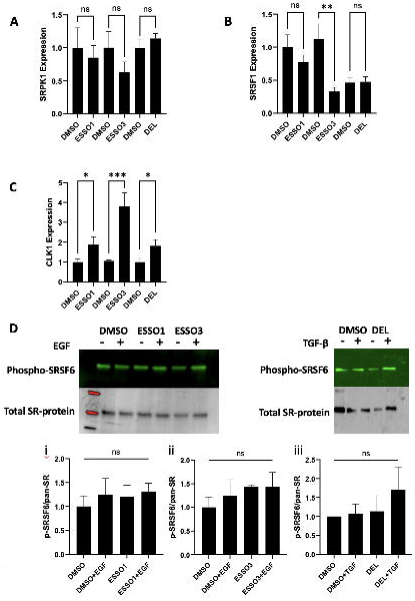
The effect of ESSO 1, ESSO3 and Delphinidine on proximal and distal splice sites regulators. **A.** RNA was extracted from podocyte cells treated with ESSO1 (10 μM), ESSO3 (10 μM) or Delphinin (10 μg/ml) for 48 hrs all in growth factor-free and serum-reduced media. RNA was made into cDNA and a qRT-PCR analysis using SRPK1 primers showed a non-significant reduction in SRPK1 expression levels for ESSO1 (mean=0.85, ±0.18 SEM, ns p=0.69, N=3 repeats), ESSO3 (mean=0.63, ±0.16 SEM, ns p=0.42, N=5 repeats) and Delphinidin (mean=1.14, ±0.07 SEM, ns p=0.39, N=4 repeats) compared to the DMSO control. Data were analysed by a two-tailed unpaired t-test. **B.** RNA was extracted from podocyte cells treated with ESSO1 (10 μM), ESSO3 (10 μM) or Delphinidin (10 μg/ml) for 48 hrs all in growth factor-free and serum-reduced media. RNA was made into cDNA. Using SRSF1 primers, the qRT-PCR showed a decreased expression of SRSF1 compared to DMSO control for ESSO1 (mean=0.77, ±0.11 SEM, ns p=0.29, N= 7 repeats), ESSO3 (mean=0.329, ±0.0641 SEM, ** p=0.007, N=5 repeats) and Delphinidin (mean=0.47, ±0.08 SEM, ns p=0.37, N=4 repeats). Data were analysed by a two-tailed unpaired t-test. **C.** RNA was extracted from podocyte cells treated with ESSO1 (10 μM), ESSO3 (10 μM) or Delphinidin (10 μg/ml) for 48 hrs all in growth factor free and serum-reduced media. RNA was made into cDNA and a qRT-PCR quantification using CLK-1 primers showed a significant increase of CLK-1 expression levels for ESSO1 (mean=1.54, ±0.14 SEM, *p=0.03, N=7 repeats), ESSO3 (mean=3.789, ±0.694 SEM, *p=0.0001, N=12 repeats) and Delphinidin (mean=1.82, ±0.31 SEM, *p=0.04, N=12 repeats) compared to the DMSO control. Data were analysed by a two-tailed unpaired t-test analysis. **D.** Podocyte cells were serum-starved for 4 hrs and treated with either DMSO, ESSO1, ESSO3 (all at 10 μM) or Delphinidin (at 10 μg/ml) for 2 hrs. SR-protein phosphorylation was stimulated with or without EGF (20ng/ml) for ESSO1 and ESSO3 and Delphinidin with or without TGF (10 ng/ml) for 30min. Proteins were extracted and phosphorylated SR-proteins were immuno-precipitated with anti-SR(1H4) and the western blot was probed with mab104 primary antibody. Western blot quantification of phosphorylated SRSF6 relative to pan SR showed an increase in phosphorylated SRSF6 for ESSO1 (mean=1.31, ±0.17 SEM, p=ns, N=3 repeats), ESSO3 (mean=1.44, ±0.30 SEM, p=ns, N=3 repeats) and Delphinidin (mean=2.01, ±0.91 SEM, p=ns, N=3 repeats). Data were analysed by ordinary one-way ANOVA with a Kruskal-Wallis post-hoc analysis.

Regarding DSS selection, the mRNA expression levels of CLK1 were found to be significantly elevated after treatment with all three compounds for 48 h compared to DMSO (**Figure 3C**). Using immunoprecipitation, the expression levels of the active form of SRSF6 – phosphorylated SRSF6 – were assessed following treatment with compounds +/- growth factor stimulation. We observed no significant difference between the treatments (**Figure 3D**).

### 3.4 Other molecules involved in ESSO1 signalling

#### JNK inhibition

JNK has been reported to be involved in *VEGF-A* AS. Specifically, JNK was reported to be involved in the selection of the DSS through SRSF2 phosphorylation in the macrophage cell line, RAW264.7 (13). ESSO1 has also been reported to activate JNK through TNFα (25). Therefore, we assessed whether ESSO1 regulates *VEGF-A* AS through JNK. Podo/TERT256 cells were treated with the the JNK inhibitor SP600125 together with ESSO1 for 48 h. Western blotting analysis showed SP600125 + ESSO1 caused a significant increase in the VEGF-A_165_b/panVEGF-A_165_ ratio compared to DMSO or SP600125 treatment alone (**Figure 4A and B**).

**Figure 4.**
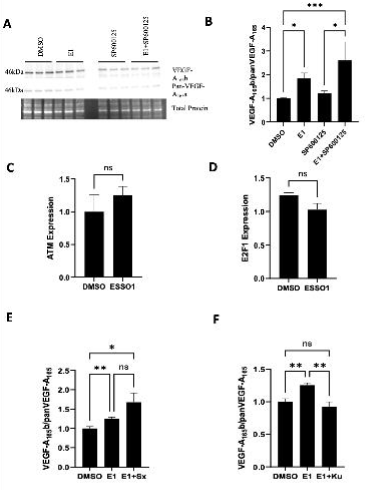
Other molecules involved in ESSO1 signalling. **A, B. JNK inhibitor SP600125 significantly increases VEGF-A165b expression with ESS01**. Proteins extracted from podocyte cells treated with ESSO1(10μM), SP600125(10μM) or ESSO1(10μM) and SP600125(10μM), for 48hours all in growth factor free and serum reduced media. Western blot performed using the proteins extracted, showed **A**. a significant increase in the ratio of VEGF-A165b/panVEGF-A165 compared to DMSO control (mean=1.841, ±0.=231 SEM, *p=0.019, N=15) and **B**. and. a further significant increase in the ratio of VEGF-A165b/panVEGF-A165 compared to control (mean=6.02, ±0.77 SEM, ***p=0.0005, N=15) **C. ESSO1 effect on ATM expression.** RNA was extracted from podocyte cells treated with ESSO1 (10 μM) for 48 hrs all in growth factor free and serum-reduced media. RNA was made into cDNA and a qRT-PCR analysis shows ATM (mean=1.25 ns p=0.23, ±0.13 SEM, N=4 repeats) expression levels are increased compared to DMSO control after ESSO1 treatment. Data were analysed by a two-tailed unpaired t-test. **D. ESSO1 treatment reduces E2F1 expression** RNA was extracted from podocyte cells treated with ESSO1 (10 μM) for 48 hrs all in growth factor free and serum-reduced media. RNA was made into cDNA and a qRT-PCR analysis shows E2F1 (mean=1.02 ns p=0.09, ±0.09 SEM, N=3 repeats) expression is reduced compared to DMSO control after ESSO1 treatment. Data were analysed by a two-tailed unpaired t-test. **E, F. ESSO1 combined with Sphinx31 (SRPK1 inhibitor) significantly increases VEGF-A165b/panVEGF-A165 expression ratio. However, ESSO1 combined with KuWaI151 (CLK-1 inhibitor) significantly reduces VEGF-A165b/panVEGF-A165 expression ratio.** **E.** Western blot quantification of VEGF-A isoform expression levels. HEK293 cells were treated with or without DMSO, ESSO1 (E1), ESSO1+ Sphinx31 (Sx) 3µM (cells pre-treated with Sphinx31 2hrs before ESSO1 treatment) or ESSO1+ KuWal151 (Ku) at 0.2µM (cells pre-treated with Kuwal 2hrs before ESSO1 treatment). Cells were incubated at 37°C for 48 hrs and proteins were extracted for a western blot. A. VEGF-A165b/panVEGF-A165 There is an overall significant difference between the treatment (**p=0.0028) and a pairwise significant increase between DMSO and E1 (*p=0.0207) and between E1 and E1+Sx (**p=0.0045) **F.** VEGF-A165b/panVEGF-A165. There is an overall significant difference between the treatment (**p=0.0027) and a pairwise significant increase between DMSO and E1 (**p=0.0038) and a significant decrease between E1 and E1+Ku (**p=0.0059)

#### ATM and E2F1

ATM is reported to be activated by ESSO1 through DNA damage by topoisomerase II alpha (TopIIα) inhibition in the macrophage cell line RAW264.7 (25). ATM is also known to target E2F1 and p38 MAPK in H460 cells and U2OS cells, respectively (26,27). E2F1 has also been implicated in DSS selection through SRSF2 in H358 cells (28). Therefore, we assessed the mRNA expression levels of ATM and E2F1 in Podo/TERT256 cells treated with ESSO1 for 48 h. We observed no significant changes in the expression of ATM and E2F1 compared to DMSO (**Figure 4C, D**).

#### SRPK1 and CLK1 inhibition

To verify if ESSO1 utilises SRPK1 and CLK1 in *VEGF-A* AS, SRPK1 was inhibited using SPHINX31 (29) and CLK1 was inhibited with KUWal151 (30), in the presence of ESSO1. We observed a significant increase in the VEGF-A_165_b/panVEGF-A_165_ ratio when cells were treated with SPHINX31 + ESSO1 (**Figure 4E**). Furthermore, KuWal151 + ESSO1 caused a significant decrease in the VEGF-A_165_b/panVEGF-A_165_ ratio compared to ESSO1 treatment alone (**Figure 4F**).

### 3.5 Molecules involved in ESSO3 signalling

#### hnRNPA1, hnRNPH, and Sam68

ESSO3 is known to regulate the expression of splicing factors, including hnRNPA1, hnRNPH, and Sam68, via the inhibition of c-MYC (14). Furthermore, these splicing factors regulate other splicing factors involved in VEGF-A AS. hnRNPA1 and Sam68 are known to antagonise and regulate SRSF1, respectively (31,32). Meanwhile, hnRNPH is known to affect p38 MAPK activation (33). qRT-PCR was performed to verify how ESSO3 affects the expression of these splicing factors in podocyte cells. Following treatment with ESSO3, there was no significant change in the expression of hnRNPA1 and Sam68 compared to DMSO (**Figure 5A, C**). In contrast, hnRNPH expression was significantly increased with ESSO3 compared to DMSO (**Figure 5B**).

**Figure 5.**
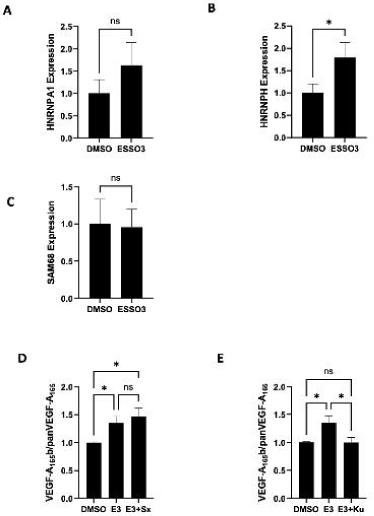
Other molecules involved in ESSO3 signalling. **A. ESSO3 treatment increases HNRNPA1 expression** RNA was extracted from podocyte cells treated with ESSO3 (10 μM) for 48 hrs all in growth factor free and serum-reduced media. RNA was made into cDNA and a qRT-PCR analysis shows HNRNPA1 (mean=1.63 ns p=0.30, ±0.51 SEM, N=11 repeats) expression levels are increased compared to DMSO control after ESSO3 treatment. Data were analysed by a two-tailed lognormal unpaired t-test. **B.** ESSO3 treatment significantly upregulates HNRNPH expression. RNA was extracted from podocyte cells treated with ESSO3 (10 μM) for 48 hrs all in growth factor free and serum-reduced media. RNA was made into cDNA and a qRT-PCR analysis shows HNRNPH (mean=1.79, *p=0.024, ±0.34 SEM, N=11 repeats) expression levels are increased compared to DMSO control after ESSO3 treatment. Data were analysed by a two-tailed lognormal unpaired t-test. **C.** ESSO3 does not affect SAM68 expression. RNA was extracted from podocyte cells treated with ESSO3 (10 μM) for 48 hrs all in growth factor free and serum-reduced media. RNA was made into cDNA and a qRT-PCR analysis shows SAM68 (mean=0.95 ns p=0.91, ±0.24 SEM, N=11 repeats) expression levels are increased compared to DMSO control after ESSO3 treatment. Data were analysed by a two-tailed lognormal unpaired t-test. **D.** Western blot quantification of VEGF-A isoform expression levels. HEK293 cells were treated with or without DMSO, ESSO3 (E3), ESSO3+ Sphinx31(Sx) 3 µM (cells pre-treated with Sphinx31 2hrs before ESSO3 treatment) or ESSO3+ KuWal151(Ku) at 0.2 µM (cells pre-treated with Kuwal 2hrs before ESSO3 treatment). Cells were incubated at 37°C for 48 hrs and proteins were extracted for a western blot. A. VEGF-A_165_b/panVEGF-A_165_. There is an overall significant difference between the treatment (**p=0.0083) and a pairwise significant increase between DMSO and E3 (*p=0.0191) and between DMSO and E3+Sx (*p=0.0292) **E.** VEGF-A_165_b/panVEGF-A_165_. There is an overall significant difference between the treatment (**p=0.0085) and a pairwise significant increase between DMSO and E3 (*p=0.0191) and a significant decrease between E3 and E3+Ku (*p=0.0455)

#### SRPK1 and CLK1 inhibition

We aimed to determine whether ESSO3 utilises SRPK1 and CLK1 in regulating VEGF-A AS in podocytes. SRPK1 was inhibited using Sphinx31 (34) and CLK1 was inhibited with KUWal151 (35) in the presence of ESSO3. The VEGF-A_165_b/panVEGF-A_165_ ratio was significantly increased when cells were treated with Sphinx31+ESSO3 compared to DMSO (**Figure 5D**). Furthermore, KuWal151+ESSO3 treatment significantly increased the VEGF-A_165_b/panVEGF-A_165_ratio compared to ESSO3 treatment alone (**Figure 5E**), indicating that ESSO3 regulates VEGF-A splicing through CLK1.

### 3.6 Molecules involved in Delphinidin signalling

#### AMPK and p32

Delphinidin has been shown to activate AMPK and phospho-AMPK in MDA-MB-453 cells and BT474 cells (36). AMPK is known to regulate the expression of SRSF1 through p32 in HeLa cells, as well as activate p38 MAPK phosphorylation in rat heart papillary muscles (37,38). A qRT-PCR was performed to assess the expression of AMPK and P32 in podocytes after delphinidin treatment. AMPK expression was significantly increased following delphinidin treatment compared to the DMSO control (**Figure 6A**). p32 expression was not significantly changed (**Figure 6B**).

**Figure 6.**
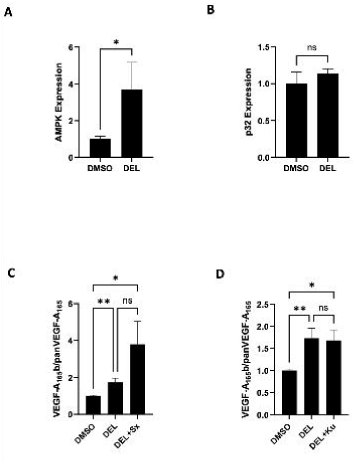
Molecules involved in Delphinidine signaling. **A. AMPK expression level is increased after treatment with Delphinidin.** RNA was extracted from podocyte cells treated with Delphinin (10 μg/ml) for 48 hrs all in growth factor free and serum-reduced media. RNA was made into cDNA and a qRT-PCR analysis AMPK (*p=0.0497, N=6 repeats) expression is increased compared to DMSO control after Delphinidin treatment. Data were analysed by a non-parametric Kolmogorov-Smirnov test. **B.** p32 expression level is not significantly changed after treatment with Delphinidin. RNA was extracted from podocyte cells treated with Delphinidin (10 μg/ml) for 48 hrs all in growth factor-free and serum-reduced media. RNA was made into cDNA and a qRT-PCR analysis shows p32 (mean=1.14 ns p=0.44, ±0.06 SEM, N=4 repeats) expression is increased compared to DMSO control after Delphinidin treatment. Data were analysed by a two-tailed unpaired t-test. **C.** Western blot quantification of VEGF-A isoform expression levels. HEK293 cells were treated with or without DMSO, Delphinidin, Delphinidin+ Sphinx31 3µM (cells pre-treated with Sphinx31 2hrs before DEL treatment) or Delphinidin+ KuWal151(Ku) at 0.2µM (cells pre-treated with Kuwal 2hrs before DEL treatment). Cells were incubated at 37°C for 48 hrs and proteins were extracted for a western blot. A. VEGF-A165b/panVEGF-A165 There is an overall significant difference between the treatment (***p=0.0001) and a pairwise significant increase between DMSO and DEL (**p=0.0066) and between DMSO and DEL+Sx (*p=0.020) **D.** VEGF-A_165_b/panVEGF-A_165_. There is an overall significant difference between the treatment (**p=0.0025) and a pairwise significant increase between DMSO and DEL (**p=0.0066) and between DMSO and DEL+Ku (*p=0.0317)

#### SRPK1 and CLK1 inhibition

To verify whether delphinidin utilises SRPK1 and CLK1 in VEGF-A AS, SRPK1 was inhibited using Sphinx31 (34) and CLK1 was inhibited with KUWal151 (35) in the presence of delphinidin. Sphinx31+delphinidin resulted in a significant increase in the VEGF-A_165_b/panVEGF-A_165_ ratio compared to DMSO (**Figure 6C**). Furthermore, KuWal151+delphinidin significantly increased the VEGF-A_165_b/panVEGF-A_165_ expression ratio compared to DMSO, but there was no significant difference compared to delphinidin alone (**Figure 6D**).

## Discussion

### ESSO1 (Trovafloxacin (TVX)) signalling regulation of VEGF-A splicing

ESSO1 has previously been shown to modulate VEGF-A alternative splicing (AS) in PC3 cells (14). Here, we demonstrate that ESSO1 also regulates VEGF-A AS in podocytes (**Figure 1**) and try to elucidate its mechanism (**Figure 7A**). ESSO1 did not affect SRPK1 and SRSF1 expression levels. However, ESSO1 significantly upregulated CLK1 expression but not phosphorylated SRSF6. Since CLK1 influences VEGF-A splice site selection, ESSO1 likely promotes VEGF-A165b/panVEGF-A165 through CLK1.

**Figure 7.**
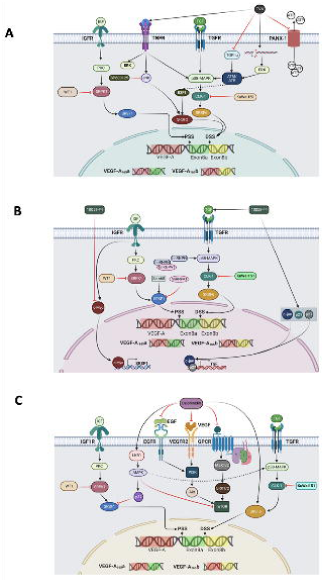
Mechanistic diagrams of the three compounds signalling to VEGF-A splicing. Proposed pathways through which ESSO1 (**A**), ESSO3 (**B**) and Delphinidine (**C**) may modulate VEGF-A AS.

From **Figure 7A**, inhibiting SRPK1 should increase the VEGF-A165b/panVEGF-A165 ratio, while inhibiting CLK1 should reduce it. Consistent with this, ESSO1 combined with SRPK1 inhibitor Sphinx31 significantly increased the ratio, whereas ESSO1 with CLK1 inhibitor KuWaI151 reduced it, supporting ESSO1’s role in signalling through the CLK1.

ESSO1 targets PANX1, TopIIα, and TNFR(25,39,40), which may indirectly affect VEGF-A AS. ESSO1 inhibits TopIIα and activates ERK via DNA damage, both of which activate ATM/ATR kinases upstream of p38 MAPK and CLK1 (25,27). qRT-PCR revealed no significant changes in ATM expression after ESSO1 treatment. Similarly, E2F1, a transcription factor linked to VEGF-A AS via SRSF2 and regulated by ATM (26,28), showed no significant change.

Further, ESSO1 has been reported to activate JNK, p38 MAPK, and ERK through TNFR (25). JNK modulates VEGF-A AS in macrophages via SRSF2 (13). To test JNK involvement in podocytes, we used SP600125 to inhibit JNK during ESSO1 treatment. Surprisingly, VEGF-A165b/panVEGF-A165 ratio increased threefold, suggesting ESSO1 acts independently of JNK and that SP600125 may enhance AS via p38 MAPK phosphorylation (41). Thus, ESSO1 and SP600125 may synergistically upregulate VEGF-A165b through p38 MAPK.

### ESSO3 (5-[(4-Ethylphenyl) methylene]-2-2thioxo-4-thiozolidonone (10058-F4)) regulation of VEGF-A splicing

We have shown ESSO3 regulates VEGF-A alternative splicing (AS) in podocytes (**Figure 1B**) and has previously been shown to modulate VEGF-A AS in PC3 cells (14). Our results (**Figure 7B**) show that ESSO3 significantly reduces nephrin expression while SRPK1 remains largely unchanged. The inhibition of c-Myc activity is likely to impact diverse transcriptional pathways (47), including those regulating nephrin. This highlights the importance of further investigation. As ESSO3 is a c-MYC inhibitor and c-MYC promotes SRSF1 transcription (42), its inhibition should explain the observed reduction in SRSF1. Conversely, CLK1, a key DSS regulator, was significantly upregulated, suggesting ESSO3 increases the VEGF-A165b/panVEGF-A165 ratio via the CLK1. However, ESSO3 did not affect phosphorylated SRSF6 expression.

To confirm CLK1 involvement in VEGF-A AS, we combined ESSO3 with SRPK1 or CLK1 inhibitor. ESSO3 plus Sphinx31 significantly increased the VEGF-A165b/panVEGF-A165 ratio, supporting the use of the DSS when SRPK1 is inhibited. In contrast, ESSO3 plus KuWaI151 reduced the ratio compared to ESSO3 alone, confirming ESSO3 acts primarily through CLK1.

ESSO3 has been reported to upregulate c-Jun and cell cycle regulators p21 and p27 (18). Since c-Jun interacts with p27 to co-activate TGFβ2 transcription (43)Increased TGFβ signalling may enhance TGFR activity and activate CLK1 through p38 MAPK. Given c-MYC’s role in regulating splice factors such as hnRNPA1, hnRNPH, and Sam68 (44–46)We assessed their expression post-ESSO3 treatment. hnRNPA1 and Sam68 showed no significant changes, while hnRNPH was significantly upregulated. This is unexpected for a c-MYC inhibitor; however, it may indicate a compensatory increase due to stress (48) or the use of alternative pathways. hnRNPH can enhance NF-κB, p38 MAPK, and JNK phosphorylation (33), which may influence DSS selection since p38 MAPK is known to phosphorylate CLK1.

### Delphinidin regulation of VEGF-A splicing

Delphinidin, an anthocyanin abundant in DIAVIT, modulates VEGF-A alternative splicing (AS) in podocytes (15). Our findings (**Figure 7C**) show that SRPK1 and SRSF1 expression were unchanged after delphinidin treatment, while CLK1, a key DSS regulator, was significantly upregulated. Given that previous studies reported a significant increase in p-SRSF6 at 1 hour (15)Unsurprisingly, we observed only a non-significant increase at 30 minutes, as the shorter stimulation period may not have been sufficient for a significant effect

To confirm splice site usage, SRPK1 and CLK1 were inhibited during delphinidin treatment. Sphinx31 combined with delphinidin significantly increased the VEGF-A165b/panVEGF-A165 ratio, indicating SRPK1 inhibition favours DSS usage. KuWal151 with delphinidin showed no significant change compared to delphinidin alone but increased compared to DMSO, suggesting delphinidin primarily acts via CLK1 while potentially engaging additional pathways, possibly directly activating SRSF6.

Delphinidin targets multiple pathways, notably inhibiting mTOR via VEGFR2 and EGFR suppress (49,50,54) and activating AMPK through LKB1 (36). AMPK can downregulate SRSF1 by activating p32, stabilising hnRNPA1 (an SRSF1 antagonist), and suppressing c-MYC or IGF-1 (37,42,52,55). Consistent with this, AMPK expression increased significantly, while p32 expression was not affected. AMPK activation may also promote p38 MAPK phosphorylation, which activates CLK1 for DSS selection (38).

mauni. Evidence from two independent RNA-seq analyses of human glomerular tissue indicated CLK1 was differentially expressed, with both studies demonstrating reduced CLK1 expression in patients with DN relative to healthy volunteers (55,56) – see **Figure 8** and **Table 3**.

**Figure 8.**
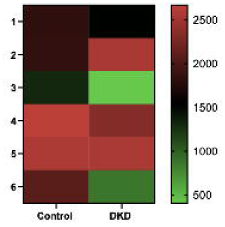
CLK1 is downregulated in in DN glomerulus. Part of a heat map and dendrogram of unsupervised hierarchical clustering based on similarity in gene-expression patterns of the six different arrays. The different colours showed the normalized Z-score for each gene. Green are downregulated genes and red are upregulated genes.

While most research in cancer, ageing, and Alzheimer’s disease highlights the therapeutic potential of CLK1 inhibition (57–59), there is also evidence that enhancing CLK1 activity may be advantageous in some disease state. For instance, it has been shown that upregulating CLK1 could be beneficial in diseases such as frontotemporal dementia and other tauopathies. The promotion of CLK1 in skipping tau exon 10 restored the normal 3R/4R tau ratio. Since excess 4R tau drives aggregation and neurodegeneration, increasing CLK1 activity corrected miss-splicing(60). Similarly, modulating CLK1 activity to restore its function presents a therapeutic potential for future research in DN.

Nevertheless, given that sustained or systemic elevation of CLK1 activity could promote cell proliferation and oncogenic splicing patterns, any therapeutic approach to increase CLK1 function should be highly targeted and context-specific, with the aim of restoring physiological balance.

## Acknowledgements

Funding for this study was supported by Diabetes UK, Richard Bright VEGF Research Trust and BBSRC grant BB/J007293/2

Many thanks to Jake Hunter, James Wright and Yihuan Liu for assistance with some of the experiments.

## Notes

### Competing Interest Statement

The authors have declared no competing interest.

## References

1. Black DL. MECHANISMS OF ALTERNATIVE PRE-MESSENGER RNA SPLICING. 2003 [cited 2020 Apr 9]; Available from: www.annualreviews.org

2. Graveley BR. Alternative splicing: increasing diversity in the proteomic world. Trends in Genetics. 2001 Feb 1;17(2):100–7.

3. Wang ET, Sandberg R, Luo S, Khrebtukova I, Zhang L, Mayr C, et al. Alternative isoform regulation in human tissue transcriptomes. Nature. 2008 Nov 27;456(7221):470–6.

4. Yang X, Coulombe-Huntington J, Kang S, Sheynkman GM, Hao T, Richardson A, et al. Widespread Expansion of Protein Interaction Capabilities by Alternative Splicing. Cell. 2016 Feb 11;164(4):805–17.

5. Bates DO, Cui TG, Doughty JM, Winkler M, Sugiono M, Shields JD, et al. VEGF165b, an Inhibitory Splice Variant of Vascular Endothelial Growth Factor, Is Down-Regulated in Renal Cell Carcinoma. Cancer Res. 2002;62(14).

6. Woolard J, Wang WY, Bevan HS, Qiu Y, Morbidelli L, Pritchard-Jones RO, et al. VEGF165b, an inhibitory vascular endothelial growth factor splice variant: Mechanism of action, in vivo effect on angiogenesis and endogenous protein expression. Cancer Res. 2004 Nov 1;64(21):7822–35.

7. Nowak DG, Woolard J, Amin EM, Konopatskaya O, Saleem MA, Churchill AJ, et al. Expression of pro- and anti-angiogenic isoforms of VEGF is differentially regulated by splicing and growth factors. J Cell Sci. 2008 Oct 15;121(20):3487–95.

8. Kawamura H, Li X, Harper SJ, Bates DO, Claesson-Welsh L. Vascular endothelial growth factor (VEGF)-A165b is a weak in vitro agonist for VEGF receptor-2 due to lack of coreceptor binding and deficient regulation of kinase activity. Cancer Res. 2008 Jun 15;68(12):4683–92.

9. Bevan HS, Van Den Akker NMS, Qiu Y, Polman JAE, Foster RR, Yem J, et al. The Alternatively Spliced Anti-Angiogenic Family of VEGF Isoforms VEGFxxxb in Human Kidney Development. Nephron Physiol [Internet]. 2008 Dec [cited 2022 Nov 17];110(4):p57–67. Available from: https://www.karger.com/Article/FullText/177614

10. Oltean S, Qiu Y, Ferguson JK, Stevens M, Neal C, Russell A, et al. Vascular endothelial growth factor-A165b is protective and restores endothelial glycocalyx in diabetic nephropathy. Journal of the American Society of Nephrology. 2015 Aug 1;26(8):1889–904.

11. Wagner KD, El Maï M, Ladomery M, Belali T, Leccia N, Michiels JF, et al. Altered VEGF Splicing Isoform Balance in Tumor Endothelium Involves Activation of Splicing Factors Srpk1 and Srsf1 by the Wilms’ Tumor Suppressor Wt1. Cells 2019, Vol 8, Page 41 [Internet]. 2019 Jan 11 [cited 2022 Nov 17];8(1):41. Available from: https://www.mdpi.com/2073-4409/8/1/41/htm

12. Manetti M, Guiducci S, Romano E, Ceccarelli C, Bellando-Randone S, Conforti ML, et al. Clinical/Translational Research Overexpression of VEGF 165 b, an Inhibitory Splice Variant of Vascular Endothelial Growth Factor, Leads to Insufficient Angiogenesis in Patients With Systemic Sclerosis. 2011 [cited 2022 Nov 17]; Available from: http://circres.ahajournals.org

13. Kikuchi R, Nakamura K, MacLauchlan S, Ngo DTM, Shimizu I, Fuster JJ, et al. An antiangiogenic isoform of VEGF-A contributes to impaired vascularization in peripheral artery disease. Nature Medicine 2014 20:12 [Internet]. 2014 Nov 2 [cited 2022 Nov 17];20(12):1464–71. Available from: https://www.nature.com/articles/nm.3703

14. Star E, Stevens M, Gooding C, Smith CWJ, Li L, Ayine ML, et al. A drug-repositioning screen using splicing-sensitive fluorescent reporters identifies novel modulators of VEGF-A splicing with anti-angiogenic properties. Oncogenesis 2021 10:5 [Internet]. 2021 May 3 [cited 2022 Oct 16];10(5):1–12. Available from: https://www.nature.com/articles/s41389-021-00323-0

15. Stevens M, Neal CR, Craciun EC, Dronca M, Harper SJ, Oltean S. The natural drug DIAVIT is protective in a type II mouse model of diabetic nephropathy. PLoS One. 2019 Mar 1;14(3).

16. Brighty KE, Gootz TD. The chemistry and biological profile of trovafloxacin. J Antimicrob Chemother [Internet]. 1997 [cited 2022 Nov 18];39 Suppl B(SUPPL. B):1–14. Available from: https://pubmed.ncbi.nlm.nih.gov/9222064/

17. Lucena MI, Andrade RJ, Rodrigo L, Salmerón J, Alvarez A, Lopez-Garrido MJ, et al. Trovafloxacin-induced acute hepatitis. Clinical Infectious Diseases [Internet]. 2000 Feb 1 [cited 2022 Nov 18];30(2):400–1. Available from: https://academic.oup.com/cid/article/30/2/400/382819

18. Huang MJ, Cheng Y chih, Liu CR, Lin S, Liu HE. A small-molecule c-Myc inhibitor, 10058-F4, induces cell-cycle arrest, apoptosis, and myeloid differentiation of human acute myeloid leukemia. Exp Hematol. 2006 Nov 1;34(11):1480–9.

19. Lai D, Huang M, Zhao L, Tian Y, Li Y, Liu D, et al. Delphinidin-induced autophagy protects pancreatic β cells against apoptosis resulting from high-glucose stress via AMPK signaling pathway. Acta Biochim Biophys Sin (Shanghai) [Internet]. 2019 Dec 13 [cited 2022 Nov 18];51(12):1242–9. Available from: https://academic.oup.com/abbs/article/51/12/1242/5645125

20. Lay AC, Tran VDT, Nair V, Betin V, Hurcombe JA, Barrington AF, et al. Profiling of insulin-resistant kidney models and human biopsies reveals common and cell-type-specific mechanisms underpinning Diabetic Kidney Disease. Nat Commun [Internet]. 2024 Dec 1 [cited 2025 Nov 14];15(1). Available from: https://pubmed.ncbi.nlm.nih.gov/39562547/

21. Lay AC, Hurcombe JA, Betin VMS, Barrington F, Rollason R, Ni L, et al. Prolonged exposure of mouse and human podocytes to insulin induces insulin resistance through lysosomal and proteasomal degradation of the insulin receptor. Diabetologia [Internet]. 2017 Nov 1 [cited 2025 Nov 14];60(11):2299–311. Available from: https://pubmed.ncbi.nlm.nih.gov/28852804/

22. Gammons M V., Fedorov O, Ivison D, Du C, Clark T, Hopkins C, et al. Topical Antiangiogenic SRPK1 Inhibitors Reduce Choroidal Neovascularization in Rodent Models of Exudative AMD. Invest Ophthalmol Vis Sci [Internet]. 2013 [cited 2025 Nov 14];54(9):6052. Available from: https://pmc.ncbi.nlm.nih.gov/articles/PMC3771558/

23. Chomzynski P. Single-Step Method of RNA Isolation by Acid Guanidinium Thiocyanate–Phenol–Chloroform Extraction. Anal Biochem. 1987 Apr;162(1):156–9.

24. Zahler AM, Lane WS, Stolk JA, Roth MB. SR proteins: a conserved family of pre-mRNA splicing factors. 1992;

25. Poulsen KL, Albee RP, Ganey PE, Roth RA. Trovafloxacin Potentiation of Lipopolysaccharide-Induced Tumor Necrosis Factor Release from RAW 264.7 Cells Requires Extracellular Signal-Regulated Kinase and c-Jun N-Terminal Kinase s. THE JOURNAL OF PHARMACOLOGY AND EXPERIMENTAL THERAPEUTICS J Pharmacol Exp Ther [Internet]. 2014 [cited 2022 Nov 18];349:185–91. Available from: 10.1124/jpet.113.211276

26. Lin WC, Lin FT, Nevins JR. Selective induction of E2F1 in response to DNA damage, mediated by ATM-dependent phosphorylation. 2001 [cited 2022 Nov 18]; Available from: https://www.genesdev.org

27. Reinhardt HC, Aslanian AS, Lees JA, Yaffe MB. p53-Deficient Cells Rely on ATM- and ATR-Mediated Checkpoint Signaling through the p38MAPK/MK2 Pathway for Survival after DNA Damage. Cancer Cell. 2007 Feb 13;11(2):175–89.

28. Merdzhanova G, Gout S, Keramidas M, Edmond V, Coll JL, Brambilla C, et al. The transcription factor E2F1 and the SR protein SC35 control the ratio of pro-angiogenic versus antiangiogenic isoforms of vascular endothelial growth factor-A to inhibit neovascularization in vivo. Oncogene 2010 29:39 [Internet]. 2010 Jul 19 [cited 2022 Nov 18];29(39):5392–403. Available from: https://www.nature.com/articles/onc2010281

29. Batson J, Toop HD, Redondo C, Babaei-Jadidi R, Chaikuad A, Wearmouth SF, et al. Development of Potent, Selective SRPK1 Inhibitors as Potential Topical Therapeutics for Neovascular Eye Disease. 2017;

30. Walter A, Chaikuad A, Helmer R, Loaë N, Preu L, Ott I, et al. Molecular structures of cdc2-like kinases in complex with a new inhibitor chemotype. 2018;

31. Mayeda A, Helfman DM, Krainer AR. Modulation of exon skipping and inclusion by heterogeneous nuclear ribonucleoprotein A1 and pre-mRNA splicing factor SF2/ASF. Mol Cell Biol [Internet]. 1993 May [cited 2022 Nov 18];13(5):2993–3001. Available from: https://journals.asm.org/doi/10.1128/mcb.13.5.2993-3001.1993

32. Valacca C, Bonomi S, Buratti E, Pedrotti S, Baralle FE, Sette C, et al. Sam68 regulates EMT through alternative splicing–activated nonsense-mediated mRNA decay of the SF2/ASF proto-oncogene. Journal of Cell Biology [Internet]. 2010 Oct 4 [cited 2022 Nov 18];191(1):87–99. Available from: https://www.jcb.org/cgi/doi/10.1083/jcb.201001073JCB87

33. Sun T, Wei C, Wang D, Wang X, Wang J, Hu Y, et al. The small RNA mascRNA differentially regulates TLR-induced proinflammatory and antiviral responses. 2021 [cited 2022 Nov 18]; Available from: 10.1172/jci.

34. Batson J, Toop HD, Redondo C, Babaei-Jadidi R, Chaikuad A, Wearmouth SF, et al. Development of Potent, Selective SRPK1 Inhibitors as Potential Topical Therapeutics for Neovascular Eye Disease. 2017 [cited 2023 May 3]; Available from: https://pubs.acs.org/sharingguidelines

35. Walter A, Chaikuad A, Helmer R, Loaë N, Preu L, Ott I, et al. Molecular structures of cdc2-like kinases in complex with a new inhibitor chemotype. 2018 [cited 2023 May 3]; Available from: 10.1371/journal.pone.0196761.g001

36. Chen J, Zhu Y, Zhang W, Peng X, Zhou J, Li F, et al. Delphinidin induced protective autophagy via mTOR pathway suppression and AMPK pathway activation in HER-2 positive breast cancer cells. BMC Cancer [Internet]. 2018 Mar 27 [cited 2022 Nov 18];18(1):1–13. Available from: https://bmccancer.biomedcentral.com/articles/10.1186/s12885-018-4231-y

37. Petersen-Mahrt SK, Estmer C, Öhrmalm C, Matthews DA, Russell WC, Akusjärvi G. The splicing factor-associated protein, p32, regulates RNA splicing by inhibiting ASF/SF2 RNA binding and phosphorylation. EMBO Journal [Internet]. 1999 Feb 15 [cited 2021 Feb 24];18(4):1014–24. Available from: https://www.embopress.org/doi/full/10.1093/emboj/18.4.1014

38. Li J, Miller EJ, Ninomiya-Tsuji J, Iii RRR, Young LH. AMP-Activated Protein Kinase Activates p38 Mitogen-Activated Protein Kinase by Increasing Recruitment of p38 MAPK to TAB1 in the Ischemic Heart. 2005 [cited 2022 Nov 19]; Available from: http://circres.ahajournals.org

39. Shaw PJ, Beggs KM, Sparkenbaugh EM, Dugan CM, Ganey PE, Roth RA. Trovafloxacin Enhances TNF-Induced Inflammatory Stress and Cell Death Signaling and Reduces TNF Clearance in a Murine Model of Idiosyncratic Hepatotoxicity. Toxicological Sciences [Internet]. 2009 Oct 1 [cited 2022 Oct 16];111(2):288–301. Available from: https://academic.oup.com/toxsci/article/111/2/288/1642971

40. Poon IKH, Chiu YH, Armstrong AJ, Kinchen JM, Juncadella IJ, Bayliss DA, et al. Unexpected link between an antibiotic, pannexin channels and apoptosis. Nature 2014 507:7492 [Internet]. 2014 Mar 12 [cited 2022 Oct 17];507(7492):329–34. Available from: https://www.nature.com/articles/nature13147

41. Vaishnav D, Jambal P, Reusch JEB, Pugazhenthi S. SP600125, an inhibitor of c-jun N-terminal kinase, activates CREB by a p38 MAPK-mediated pathway. Biochem Biophys Res Commun [Internet]. 2003 [cited 2022 Nov 18];307(4):855–60. Available from: www.elsevier.com/locate/ybbrc

42. Das S, Anczuków O, Akerman M, Krainer AR. Oncogenic Splicing Factor SRSF1 Is a Critical Transcriptional Target of MYC. Cell Rep. 2012 Feb 23;1(2):110–7.

43. Yoon H, Kim M, Jang K, Shin M, Besser A, Xiao X, et al. P27 transcriptionally coregulates cJun to drive programs of tumor progression. Proc Natl Acad Sci U S A [Internet]. 2019 [cited 2022 Nov 18];116(14):7005–14. Available from: www.pnas.org/cgi/doi/10.1073/pnas.1817415116

44. Nadiminty N, Tummala R, Liu C, Lou W, Evans CP, Gao AC. NF-κB2/p52:c-Myc:hnRNPA1 pathway regulates expression of androgen receptor splice variants and enzalutamide sensitivity in prostate cancer. Mol Cancer Ther [Internet]. 2015 Aug 1 [cited 2022 Nov 18];14(8):1884–95. Available from: https://aacrjournals.org/mct/article/14/8/1884/130548/NF-B2-p52-c-Myc-hnRNPA1-Pathway-Regulates

45. Rauch J, Moran-Jones K, Albrecht V, Schwarzl T, Hunter K, Gires O, et al. c-Myc regulates RNA splicing of the A-Raf kinase and its activation of the ERK pathway. Cancer Res [Internet]. 2011 Jul 1 [cited 2022 Nov 18];71(13):4664–74. Available from: https://aacrjournals.org/cancerres/article/71/13/4664/567766/c-Myc-Regulates-RNA-Splicing-of-the-A-Raf-Kinase

46. Caggiano C, Pieraccioli M, Panzeri V, Sette C, Bielli P. c-MYC empowers transcription and productive splicing of the oncogenic splicing factor Sam68 in cancer. Nucleic Acids Res [Internet]. 2019 Jul 9 [cited 2022 Nov 18];47(12):6160–71. Available from: https://academic.oup.com/nar/article/47/12/6160/5486749

47. Sabò A, Kress TR, Pelizzola M, De Pretis S, Gorski MM, Tesi A, et al. Selective transcriptional regulation by Myc in cellular growth control and lymphomagenesis. Nature 2014 511:7510 [Internet]. 2014 Jul 9 [cited 2025 Nov 21];511(7510):488–92. Available from: https://www.nature.com/articles/nature13537

48. Wall ML, Bera A, Wong FK, Lewis SM. Cellular stress orchestrates the localization of hnRNP H to stress granules. Exp Cell Res [Internet]. 2020 Sep 1 [cited 2025 Nov 21];394(1). Available from: https://pubmed.ncbi.nlm.nih.gov/32473225/

49. Lamy S, Blanchette M, Michaud-Levesque J, Lafleur R, Durocher Y, Moghrabi A, et al. Delphinidin, a dietary anthocyanidin, inhibits vascular endothelial growth factor receptor-2 phosphorylation. Carcinogenesis [Internet]. 2006 May 1 [cited 2022 Nov 18];27(5):989–96. Available from: https://academic.oup.com/carcin/article/27/5/989/2476121

50. Meiers S, Kemény M, Weyand U, Gastpar R, Von Angerer E, Marko D. The Anthocyanidins Cyanidin and Delphinidin Are Potent Inhibitors of the Epidermal Growth-Factor Receptor. 2001 [cited 2022 Nov 18]; Available from: https://pubs.acs.org/sharingguidelines

51. Pal HC, Sharma S, Strickland LR, Agarwal J, Athar M. Delphinidin Reduces Cell Proliferation and Induces Apoptosis of Non-Small-Cell Lung Cancer Cells by Targeting EGFR/VEGFR2 Signaling Pathways. PLoS One [Internet]. 2013 [cited 2022 Nov 18];8(10):77270. Available from: www.plosone.org

52. Expert-Bezançon A, Sureau A, Durosay P, Salesse R, Groeneveld H, Lecaer JP, et al. hnRNP A1 and the SR Proteins ASF/SF2 and SC35 Have Antagonistic Functions in Splicing of β-Tropomyosin Exon 6B. Journal of Biological Chemistry. 2004 Sep 10;279(37):38249–59.

53. Ning J, Clemmons DR. AMP-Activated Protein Kinase Inhibits IGF-I Signaling and Protein Synthesis in Vascular Smooth Muscle Cells via Stimulation of Insulin Receptor Substrate 1 S794 and Tuberous Sclerosis 2 S1345 Phosphorylation. Molecular Endocrinology [Internet]. 2010 Jun 1 [cited 2022 Nov 19];24(6):1218–29. Available from: https://academic.oup.com/mend/article/24/6/1218/2706158

54. Pal HC, Sharma S, Strickland LR, Agarwal J, Athar M. Delphinidin Reduces Cell Proliferation and Induces Apoptosis of Non-Small-Cell Lung Cancer Cells by Targeting EGFR/VEGFR2 Signaling Pathways. PLoS One [Internet]. 2013 [cited 2022 Nov 18];8(10):77270. Available from: www.plosone.org

55. Baelde HJ, Eikmans M, Doran PP, Lappin DWP, De Heer E, Bruijn JA. Gene Expression Profiling in Glomeruli from Human Kidneys with Diabetic Nephropathy. American Journal of Kidney Diseases [Internet]. 2004 Apr 1 [cited 2022 Nov 19];43(4):636–50. Available from: http://www.ajkd.org/article/S0272638604000083/fulltext

56. Levin A, Reznichenko A, Witasp A, Liu P, Greasley PJ, Sorrentino A, et al. Novel insights into the disease transcriptome of human diabetic glomeruli and tubulointerstitium. Nephrology Dialysis Transplantation [Internet]. 2021 [cited 2022 Nov 19];35(12):2059–72. Available from: https://github.com/chapmanb/bcbio-nextgen

57. Jain P, Karthikeyan C, Moorthy NS, Waiker D, Jain A, Trivedi P. Human CDC2-Like Kinase 1 (CLK1): A Novel Target for Alzheimer’s Disease. Curr Drug Targets. 2014;15(5):539–50.

58. Uzor S, Porazinski SR, Li L, Clark B, Ajiro M, Iida K, et al. CDC2-like (CLK) protein kinase inhibition as a novel targeted therapeutic strategy in prostate cancer. Sci Rep [Internet]. 2021 Dec 1 [cited 2025 Jul 23];11(1):1–12. Available from: https://www.nature.com/articles/s41598-021-86908-6

59. Babu N, Pinto SM, Biswas M, Subbannayya T, Rajappa M, Mohan S V., et al. Phosphoproteomic analysis identifies CLK1 as a novel therapeutic target in gastric cancer. Gastric Cancer [Internet]. 2020 Sep 1 [cited 2025 Jul 23];23(5):796–810. Available from: https://link.springer.com/article/10.1007/s10120-020-01062-8

60. Hartmann AM, Rujescu D, Giannakouros T, Nikolakaki E, Goedert M, Mandelkow EM, et al. Regulation of Alternative Splicing of Human Tau Exon 10 by Phosphorylation of Splicing Factors. Molecular and Cellular Neuroscience [Internet]. 2001 Jul 1 [cited 2025 Nov 22];18(1):80–90. Available from: https://www.sciencedirect.com/science/article/pii/S1044743101910000

