## Supplementary material for "Mechanistic analysis of compounds that modulate VEGF-A splicing in podocytes with therapeutic potential for diabetic nephropathy": Table 1

| **Gene** | **Species** | **Primer** | **Sequence (5’→3’)** |
| --- | --- | --- | --- |
| *SRPK1* | Human | Forward | TGGCCACAGGTGACTATTTG |
|  |  | Reverse | CCCAAGGTTTCAGCTTCGT |
| *CLK1* | Human | Forward | TGAATACTATCTTGGGTTTACCGTAT |
|  |  | Reverse | CGTTTCCTGGTTTTCTGTATCATAT |
| *SRSF1* | Human | Forward | GCGGGATCAGATTACCAGGA |
|  |  | Reverse | ACTCGTGCTGAATCCTTCCA |
| *AMPK* | Human | Forward | GTCGGCACCTTCGGCAAAGTGAAG |
|  |  | Reverse | AAATTCACCATCTGACATCAT |
| *ATM* | Human | Forward | ACCTACCAAATCCCTCCACC |
|  |  | Reverse | TGTTAGATGCCAAAATGCTCC |
| *ATR* | Human | Forward | ACACATCCAGCCTAAAGAAAC |
|  |  | Reverse | TCCCCTAAACATTCCCCAC |
| *E2F1* | Human | Forward | TCACCACCACCATCATCTC |
|  |  | Reverse | CCCAAAGTCACAGTCGAAG |
| *P32* | Human | Forward | CAACATTAACAACAGCATCCC |
|  |  | Reverse | TCCTTCCATTCAGACTCGCC |
| *GAPDH* | Human | Forward | GTCTCCTCTGACTTCAACAGCG |
|  |  | Reverse | ACCACCCTGTTGCTGTAGCCAA |
| *HPRT1* | Human | Forward | CATTATGCTGAGGATTTGGAAAGG |
|  |  | Reverse | CTTGAGCACACAGAGGGCTACA |
| *VEGF-A* | Human | Forward | TTGTACAAGATCCGCAGACG |
|  |  | Reverse | ATGGATCCGTATCAGTCTTTCCTGG |

**Table 1.** List of primers used for RT-qPCR
