## Supplementary material for "Mechanistic analysis of compounds that modulate VEGF-A splicing in podocytes with therapeutic potential for diabetic nephropathy": Table 2

| **ESSO** | **Compound** | **Description** |
| --- | --- | --- |
| **1** | Trovafloxacin | Broad-spectrum antibiotic |
| **2** | Melatonin | Hormone |
| **3** | 5-[(4-Ethylphenyl)methylene]-2-2thioxo-4-thiozolidonone | C-Myc Inhibitor |
| **4** | N6-2-(4-Aminophenyl)Ethyladenosine | A3 adenosine receptor agonist |
| **5** | 8-Bromoadenosine-3',5'-cyclophospate sodium | cAMP analog |
| **6** | Fluportine Maleate | NMDA receptor antagonist |
| **7** | RepSox | THF-beta type 1 receptor antagonist |
| **8** | GW2974 | EGFR and ErbB-2 tyrosine kinase inhibitor |
| **9** | 4-(20Aminoethyl)benzenesulfonylfluoride | Serine protease inhibitor |

**Table 1.** List of nine compounds selected after the screening of the LOPAC compounds.
