## Supplementary material for "Mechanistic analysis of compounds that modulate VEGF-A splicing in podocytes with therapeutic potential for diabetic nephropathy": Table 3

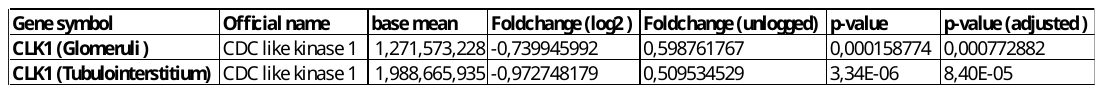

**Table 3. CLK1 is significantly differentially expressed in DN glomeruli and tubulointestitium.**

Part of a table showing DESeq2 analysis of DN [*n* = 19, age: 61 (30–85) years, chronic kidney disease stages 1–4] and living kidney donors [*n* = 20, age: 56 (30–70) years]. Benjamini–Hochberg method was used to adjust p-values for multiple testing. Genes with adjusted p<0.01 and targeted analyses with p<0.05 were considered differentially expressed
