## Supplementary Figures for "Mechanistic analysis of compounds that modulate VEGF-A splicing in podocytes with therapeutic potential for diabetic nephropathy"

### Slide 1
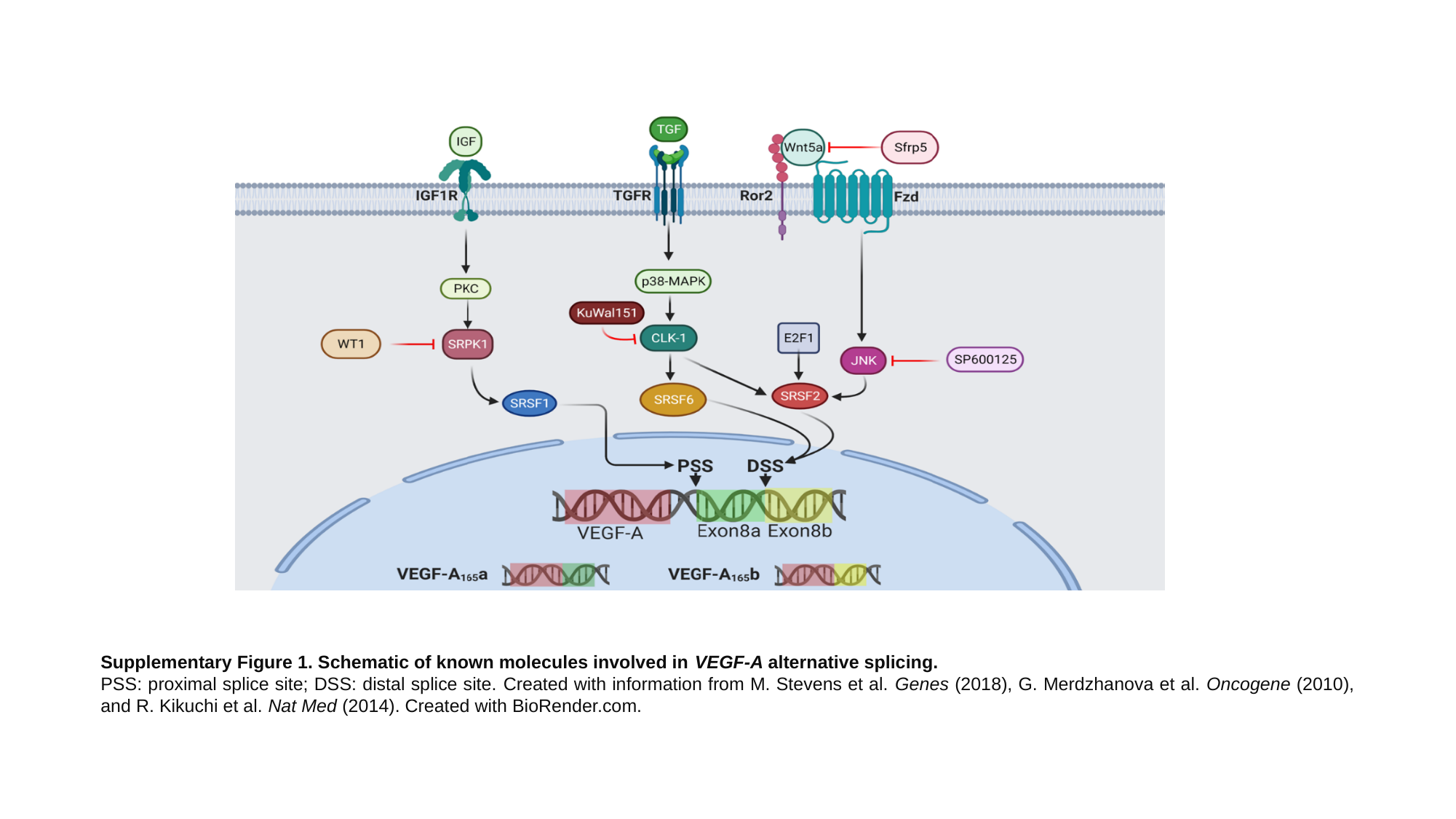

Supplementary Figure 1. Schematic of known molecules involved in VEGF-A alternative splicing.
PSS: proximal splice site; DSS: distal splice site. Created with information from M. Stevens et al. Genes (2018), G. Merdzhanova et al. Oncogene (2010), and R. Kikuchi et al. Nat Med (2014). Created with BioRender.com.
